# Computational cognitive neuroscience: Model fitting should not replace model simulation

**DOI:** 10.1101/079798

**Authors:** Stefano Palminteri, Valentin Wyart, Etienne Koechlin

## Abstract

Cognitive neuroscience, especially in the fields of learning and decision-making, is witnessing the blossoming of computational model-based analyses. Several methodological and review papers have indicated how and why candidate models should be *compared* by trading off their ability to predict the data as a function of their complexity. However, the importance of *simulating* candidate models has been so far largely overlooked, which entails several drawbacks and leads to invalid conclusions. Here we argue that the analysis of model simulations is often necessary to support the specific claims about behavioral function that most of model-based studies make. We defend this argument both informally by providing a large-scale (N>300) review of recent studies, and formally by showing how model simulations are necessary to interpret model comparison results. Finally, we propose guidelines for future work, which combine model comparison and simulation.

## Computational model comparison as theory selection

In cognitive science, and especially in the field of learning and decision-making, the utilization of computational modeling has remarkably increased during the past decade (Figure 1). Models are taking an important place also in neuroimaging and neuropsychiatry as powerful tools to understand normal, as well as diseased, cognition and brain function^1–5^. The importance of computational models in cognitive neuroscience is not surprising, since the intrinsic function of the brain is information processing at the service of goal-directed behavior, and cognitive theories can thus be formulated as computational theories^6,7^ (Box 1). As a consequence, if computational models are to be considered theories of brain function, they should be submitted to a theory selection process. In this paper we argue that the current practice of model comparison (in the purpose of selecting a winning model) often omits an important, and even necessary, step of theory selection.

**Figure 1:**
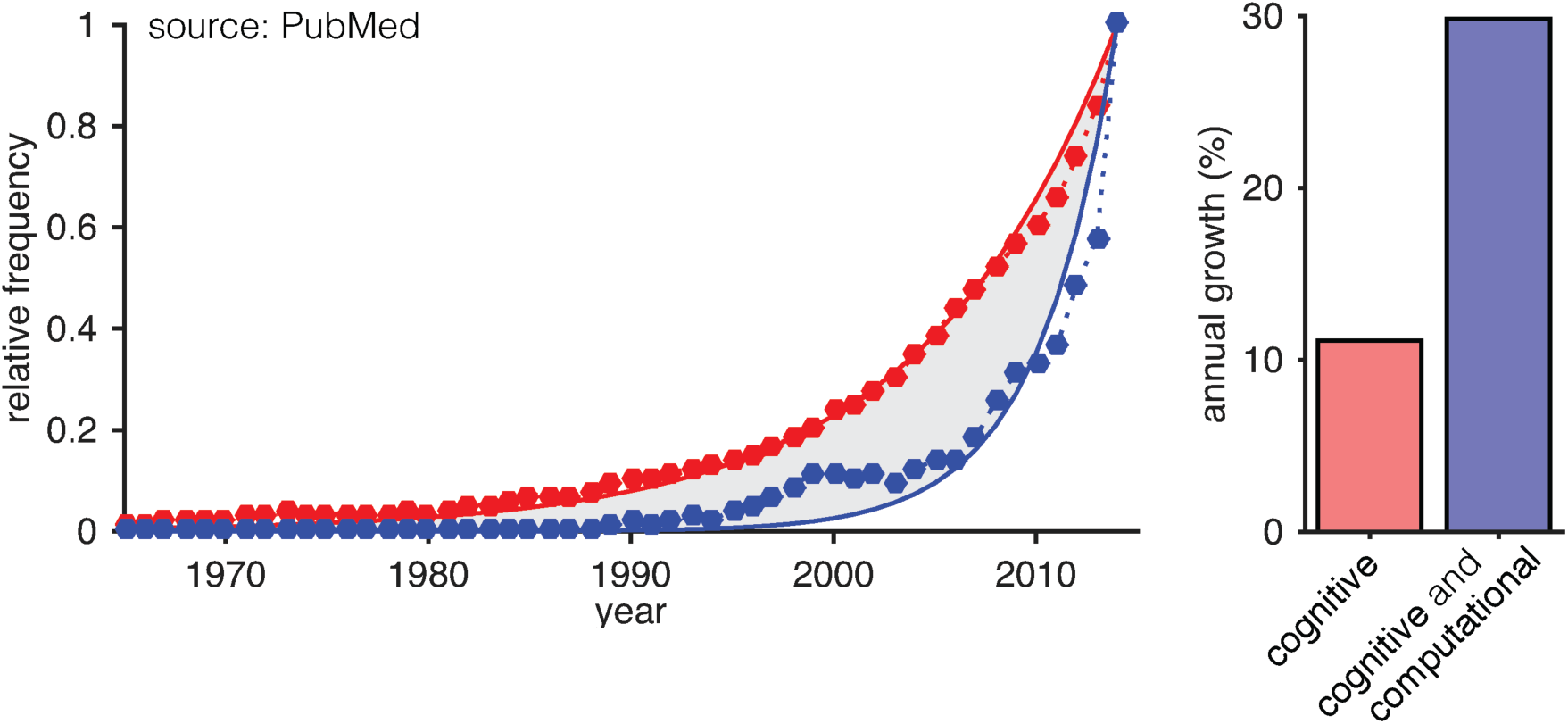
the exponential increase of computational model-based cognitive neuroscience. The curves on the left show the relative frequency of PubMed entries for “cognitive” (in red) and “cognitive and computational” (in blue) as a function of the year (from the 1970s to the 2014). Their frequencies are calculated relative to the number of entries of 2014, which are therefore defined at 1 for both curves. The bars on the left represent the estimated annual growth of the best fitting exponential curve.

#### Box 1 delineating computational modeling approaches

In cognitive neuroscience computational models can also be simply used as tools to quantify different features of the behavioral or neural activity. In this approach, the typical results consist in comparing model free parameters across conditions or group of subject^23^. In this approach computational models are not very different from statistical models, such as multiple regressions, where the statistical tests are performed on regressions coefficients, instead of parameters. The opposite view is to consider computational models as quantitative theories of the cognitive function. When a model is used as an analytical tool, model comparison is not central, because no real competition between cognitive theories is at stake.

Computational modeling can take place at several levels of description in psychology and cognitive sciences. Understanding the differences between these levels of description is mandatory, before engaging in a model comparison analysis. One important distinction is between aggregate and mechanistic models^9^. The first class of models aims at describing average (across subjects and trials) behavior in a mathematically synthetic way, such as the exponential learning equation behavior^24^. The second class of models aims at explaining how a behavior emerges on a trial-by-trial basis, such as the Rescorla-Wagner learning model ^25^. Naturally, because these two classes of models do not target the same level of description, a model selection scheme should not compare aggregate and mechanistic models. For example, in the cases presented above, an aggregate exponential learning curve could derive from a Rescorla-Wagner mechanism and both models can therefore be considered “true” once acknowledged that they do not target the same level of description. The distinction between aggregate and mechanistic models has been further developed by Marr^6^, who proposed three different levels of description for computational modeling approaches. The “computational” level deals with the goal of the computation. For example, in standard instrumental learning tasks, the goal of the computation is to maximize the occurrence of rewards. The “representational and algorithmic level” deals with how the computational theory is realized in terms of input and output (representations) and the mathematical operations in between the two (algorithms). For example, instrumental behavior is often formalized within the temporal difference-learning framework that supposes instrumental behavior driven by prediction errors^26^. Finally, the “hardware or implementational” level deals with the way the algorithm is physically implemented in the brain. For example, temporal difference reward prediction errors are supposed to be represented by phasic increase in the firing rate of the dopaminergic neurons^27^.

Again, when engaging in a model selection scheme, it is important to keep in mind which level of description is targeted. In fact, a model comparison has different meanings at the “computational”, “algorithmic” and “hardware” levels. At the “computational” level, model comparison will inform on the “goal” of the subjects and therefore their understanding of the task. Of course, this description is dissociable from the “algorithmic” level. For example, the same computational goal, such as reward maximization, could be achieved using a temporal difference or a Bayesian algorithm^28^. It also important to note that model selection at different levels may require different forms of data to be achieved. In fact, whereas at the “computational” level model selection can generally be based on aggregate behavioral data (even across subjects), the “algorithmic” level most typically requires trial-by-trial data from individual subjects. On the other side, model selection at the “implementational” level will by definition require neural recordings.

One universally recognized theory selection heuristic is Occam’s *lex parsimoniae*^a^. In short, this precept dictates that amongst “equally good” explanation of data, the less complex should be held as more likely to be true. Translated in more formal terms, there is a trade off between the complexity of a given model (which grows with its number of free parameters) and its accuracy (the likelihood of the data given the model). Different quantitative solutions have been proposed to take parsimony into account when comparing different models, based on their *predictive performance* – i.e., their ability to predict observed data^8–11^. All these methods are rooted in the Bayesian conception of statistics, based on *relative* model comparison criteria. They do not imply any absolute criterion of performance: the winning model is the model having more evidence (quality of the prediction minus model complexity) compared to a rival one^8,12^, even if all candidate models provide a poor description of important features of the fitted data.

Another important heuristic, recognized by contemporary epistemology, applies to theory selection. To propose a novel theory, the scientist should be able to show that a pre-extant theory is contradicted by an experimental observation (i.e. is falsified), whereas a novel and proposed theory is not^13^. Theory rejection represents the logic substratum of hypothesis testing as implemented in classical ‘frequentist’ statistics^12,14^. Translated into computational terms, to reject a model one rests on showing that it is not able to account for a least one behavioral (or neural) phenomenon of interest – called a *rejection criterion*. The ability of a model to reproduce the phenomenon – called the *generative performance* of the model – should be assessed by simulating the model and comparing the simulations to the observed phenomenon.

Relative model comparison criteria (i.e. various approximations of the model evidence, such as BIC, AIC,) are not appropriate to falsify models because they do not capture certain features of the fitted data: 1) they focus on the evidence *in favor* of the best, instead of evidence *against* the rival model, and 2) they are blind to the capacity of tested models to reproduce (or not) any particular phenomenon of interest. Importantly, good predictive performance do not imply good generative performance (Box 2).

#### Box 2 good predictive performance does not imply good generative performance

Typical relative model comparison criteria evaluate the predictive performance of a model, i.e. the likelihood of observing the experimental data given the model. However, a computational model can display relatively good predictive performance even if its generative performance – obtained through independent model simulations, is inaccurate. This discrepancy is well captured by a simple example, originally reported by Corrado et al^9^. Concerning weather prediction, a quite good predictive model is “tomorrow’s weather will be as today’s”. Indeed, given the important time autocorrelation of weather condition, such a simple model will provide above chance predictions. Nonetheless, it clearly appears that this model does not illuminate at all the mechanisms that govern the weather and indeed it will not allow simulating its evolution in the mid-or long-run. Unfortunately, often such temporal autocorrelation also applies to behavioral data, especially in the field of reinforcement learning (good choices naturally tend to be repeated) and decision-making (in the form of choice hysteresis^29^). To illustrate this point we simulated a win-stay lose-shift (WSLS) equivalent e of an RL model (i.e. a Q-learning with a learning rate (α=1) and a non-negative choice temperature (β)). We simulated the WSLS behavior in a deterministic two-armed bandit task in which two responses are associated with a deterministic probability of being rewarded or not, and in which the contingencies are reversed after an unknown number of choices. As expected the WSLS model is capable of adapting its responses as a function of task’s demands (i.e. it mostly chooses the “Right option” when it is rewarded and mostly chooses the “Left” option, when the contingencies switches (Figure Box 1A)). We fitted to the WSLS model an equivalent of the “tomorrow’s weather will be as today’s” model for weather prediction, adapted to this reversal task. The “Repetition” model basically instantiates choice hysteresis by assuming at each trial Q_t+1_(chosen option) = 1 and Q_t+1_(unchosen option) = 0. We fitted the “Repetition” model the virtual data to find the likelihood maximizing free parameter (β). We then plotted the trial-by-trial model estimate of choice probability of the “Repetition” model, based on the individual history of choices of the WSLS models (black dots, predictive simulations). Unsurprisingly, these predictive simulations show that the “Repetition” model is capable of switching responses after reversal and roughly follow the behavior of the WSLS model, even if the “Repetition” model is completely blind to the outcome history, which determines the WSLS behavior. However, when we simulate the “Repetition model” behavior *ex novo* in the very same task, the probability of choosing right or left remains around chance, thus revealing the model’s inner inability to follow reward contingencies and their reversal (white dots; true generative performances). This rather extreme example illustrates to which extent predictive performance, on which model comparison is performed, may be dissociated from generative performance and can lead to erroneous conclusions concerning a model’s ability to “explain” the data.

Importantly, the way in which the relative model comparison criteria are calculated may lead to an inflated representation of a model performance also in a more subtle way. The relative model comparison criteria are typically calculated based on the maximum likelihood. The calculation of the maximum likelihood requires identifying the model parameters that maximize the chance of observing the data. This imply that if an effect is present in the data, even if this effect can be, in principle, explained by the model, the likelihood maximization procedure will favor the set of free parameters that, by chance, maximize the probability of observing this effect: this closely resembles the problem that in neuroimaging is referred to as “double-dipping”. To illustrate this point we simulated a “Metalearning” model over two learning sessions of a probabilistic two-armed bandit task, in which, in each session, the model has to find out which is the most rewarding option. The options are reversed from a session to another. The “Metalearning” model has the peculiarity of using different sets of parameters (learning rate α), such as in the second session its correct response improves, by augmenting the learning rate and reducing the exploration (as if the model had integrated the task statistics to adjust its parameters to perform better; Figure Box 2B). We fitted to these virtual data a Q-learning model, whose parameters remain the same in the first and in the second session. As we have done before for the “Repetition” model, we plotted the predictive performance (i.e. the probability of choosing the most rewarding option given the subject history of choices and outcomes) and we observe a smaller, but significant, improvement of correct choice rate moving from the first to the second session, even if the parameters remain the same and there is not computational process in the Q-learning model that could underpin this effect (white dots). As a matter of fact, this effect simply arises because the meta-learning effect is present in the data and the likelihood maximization procedure has therefore favored the parameters’ values that maximize the probability of observing this effect. Of course this result does not imply at all that the Q-learning model has the ability of improving performance from one session to another. In fact, when simulating the same Q-learning models (with the parameters set retrieved with the model fitting procedure) in a new task (i.e. not giving the model the precise history of choices and outcomes of the “Metalearning” virtual subjects) we found that the Q-learning predicts no difference between the first and the second session (black dots).

To summarize, this Box draws the attention to the difference between a model’s predictive performance (on which the relative model comparison criteria are based) and generative performance. More particularly this Box illustrates how, either because of the data autocorrelation, or because of the likelihood-maximization procedure itself, model predictions resulting from model fitting often provide an inflated image of the model’s ability to account for the data, whereas only ex novo model simulations (i.e. not based on the subject history of choices and outcomes) provide an unbiased image of models’ behavioral abilities.

**Figure Box 2:**
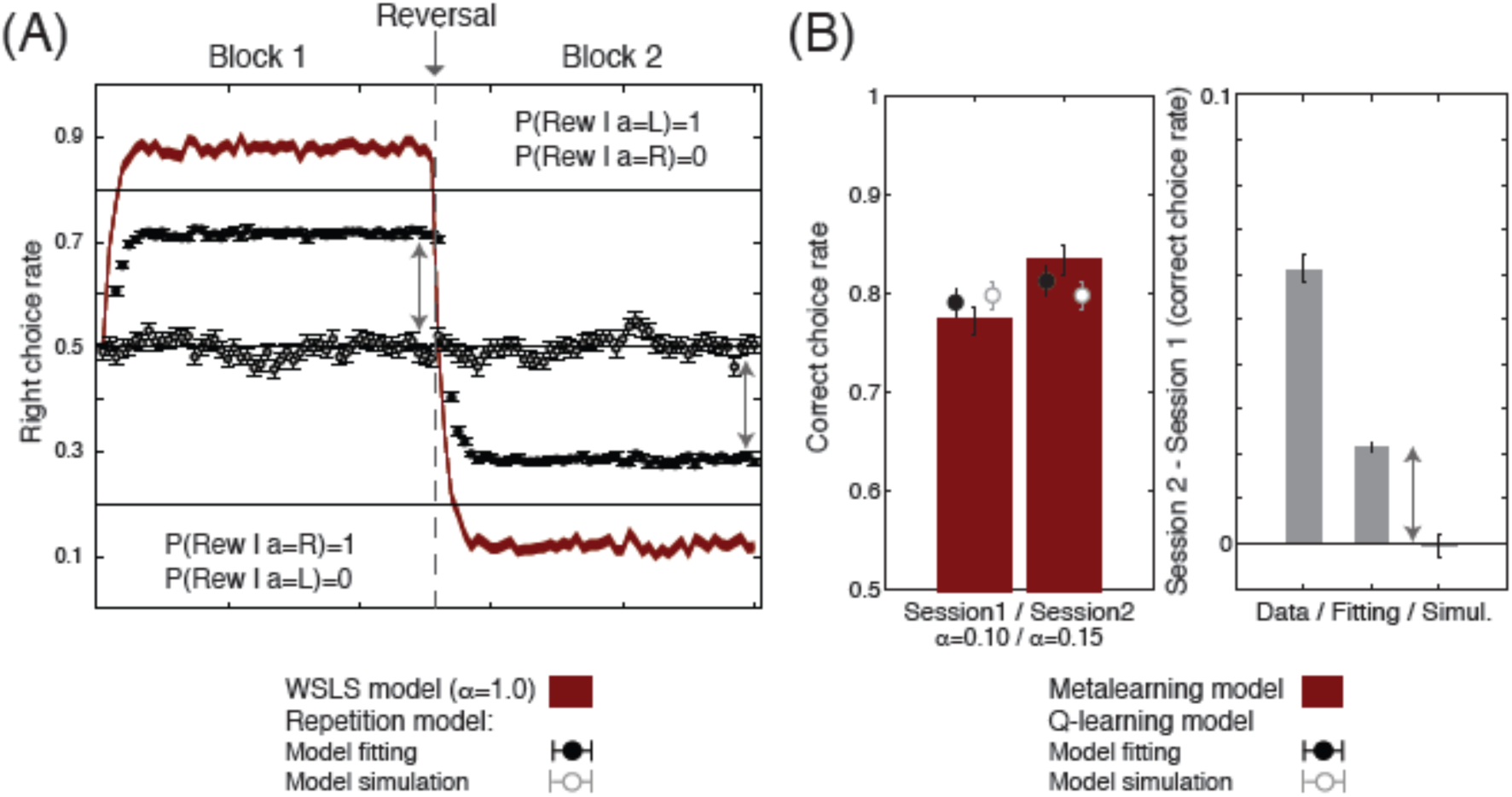
predictive vs. generative performance. **(A)** The dark red curve represents the trial-by-trial average probability (± s.e.m.) of choosing the “Right” choice in a virtual deterministic two-armed bandit task simulated with a reinforcement learning equivalent of a win-stay lose-shift model (WSLS; i.e. a Q-learning model with α=1). The black dots represent the average probability (± s.e.m.) of observing a “Right” choice, estimated by a Repetition model (i.e. a model whose Q-values update rule consist in Q(chosen option) = 1 and Q(unchosen option) = 0), based on the individual best fitting parameters and histories of choices and outcomes ((predictive performance: P(data|model, history of choices and outcomes)). The white dots represent the average probability (± s.e.m.) of observing a “Right” choice, generated by a Repetition model simulated *ex novo* on the same task (true generative performance). The dashed line represents the point in which the contingencies are reversed. The two-headed arrow highlights the discrepancy between the model fitting (predictive performance) and the model simulations (generative performance) **(B)** The leftmost panel shows the mean probability (± s.e.m.) of choosing the “Correct” choice (i.e. the most rewarding one) in a virtual probabilistic two-armed bandit task simulated with a "Metalearning model that assumes that the subject adaptively modify the free-parameters (the Metalearning model increases α and reduces β: ?α=0.05) to improve performance in the second learning session (i.e. after the dashed line, when the instrumental cues have been changed). The black dots represent the average probability (± s.e.m.) of observing a “Correct” choice, estimated by a basic Q-learning model (i.e. a model whose α and β are not modulated from the first to the second session: ?α=?β=0.0), based on the individual best fitting parameters and histories of choices and outcomes (predictive performance: P(data|model, history of choices and outcomes)). The white dots represent the average probability (± s.e.m.) of observing a “Correct” choice, generated by a Q-learning model simulated *ex novo* on the same task (true generative performance). The two-headed arrow highlights the discrepancy between the model fitting (predictive performance) and the model simulations (generative performance).

## Current common practice and associated problems

Whereas relative model comparison is now often implemented in computational modeling studies of brain activity and behavior, model simulations are only rarely performed and studied. This claim is supported by a meta-analysis of >300 studies published since 2009 in six high rank journals in the field of learning and decision-making^b^ (Science, Nature, Nature Neuroscience, Neuron, PNAS, PLOS Biology and The Journal of Neuroscience; Table 1). First, we note that in this sample, about 40% of studies (N=140) implicated computational-model based analyses. This result is in line with the observation of a remarkable increase of computational model-based analyses (Figure 1). Amongst the model-based studies, more than 50% (N=80) implemented a model selection step, based on a relative model comparison criterion. Crucially, within the pool of the model selection studies, only few (less than 20%; N=15) simulated the best as well rival/losing model(s). Here, we argue that the lack of model simulations is problematic in respect to the specific claims that are defended in most computational model-based cognitive studies.

**Table 1:** percentage of studies as a function of their model-based analytical procedure. “Computational Model” = studies report a computational model-based analysis. “Relative comparison” = studies report a model selection step implicating relative model comparison criteria, such as AIC, BIC or similar. “Best simulation” = studies report the model simulation of the best model, according to relative model selection. “Rival simulation” = studies report the simulation of the best and the rival(s) models (note that the presence of rival model simulation represents no guarantee that any statistical analysis is then performed to quantitatively assess the “similarity” of the model simulations to the actual data). The complete list of the studies included in the meta-analysis is available online).

| Sample | Computational model | Relative comparison | Best simulation | Rival simulation |
| --- | --- | --- | --- | --- |
| All studies (N=301) | 47.7% (N=143) | 27.3% (N=82) | 21.0% (N=63) | 5.7% (N=17) |
| Model studies (N=143) | - | 57.35% (N=82) | 44.1% (N=63) | 11.3% (N=17) |
| Relative studies (N=82) | - | - | 43.9% (N=36) | 18.1% (N=15) |

Typically, a modeling study proceeds along the following scheme. First, an experimental intervention (a task) is designed to elicit detectable behavioral (or neural) effects of interest. Let us consider the simplest case in which the behavioral effect of interest takes the form of different behavior in condition ‘A’ compared to ‘B’ (Figure 2). This dependent variable could be represented by very diverse observables, such as the proportion of certain choices in a decision-making task. For instance, the two conditions could be represented by different levels of reward magnitude (10 cents versus 1 euro) in an incentive motivation task, of proposed delay in a temporal discounting economic task (present versus future), or coherence in a random-dot motion discrimination task (high versus low)^15,16^. The eventual aim is then to decide which model (noted as Model 1 and Model 2) accounts for this effect, under the (more or less implicit) assumption that this model selection analysis will allow arbitrating between competing hypotheses of cognitive function. Thus, in this case, a model not being able to reproduce the behavioral difference between the two conditions A & B represents the model *rejection criterion*, in such ‘falsification’ framework.

**Figure 2:**
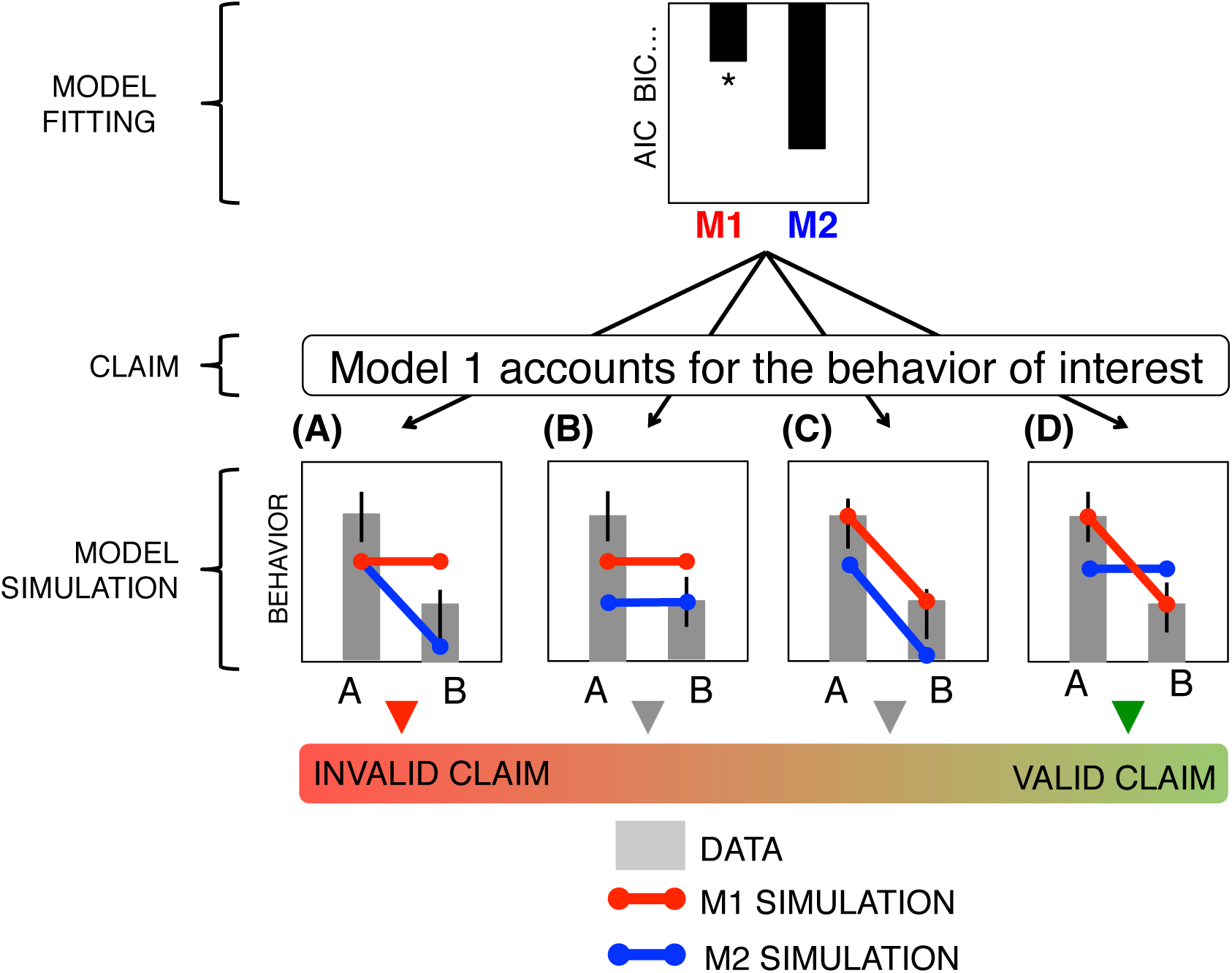
possible relationships between relative model comparison and model simulation comparison. A stereotyped case in which subjects’ behavioral data can be project into two orthogonal dimensions: the average level of performance across both conditions ((condition A+ condition B)/2) and the difference in performance between the two conditions (condition A - condition B). In all cases Model 1 is the “best” model, based on relative model comparison criteria. (**A**) Model 1 accounts for the average level of performance, but is not able to capture the behavior of interest. Model 2 is not able to explain neither the behavior of interest nor the average level of performance (**B**) Model 1 accounts for the average level of performance, but does not capture the behavior of interest. Model 2 is not able to explain neither the average level of performance nor the behavior of interest. (**C**) Model 1 accounts both the average level of performance and the behavior of interest. Model 2 is not able to explain the average level of performance, but captures the behavior of interest. (**D**) Model 1 accounts for both the average level of performance and the behavior of interest. Model 2 capture the average level of performance, but is not able to capture the behavior of interest. In the literature sample considered here only the 17% of the model comparison studies, includes the simulation of the “best” as well as the “rival(s)” models.

However, most studies (N=60; in our sample) start by determining, for each model, the free parameters that maximize the likelihood of the data given the model (model fitting). The likelihood is subsequently used to calculate a relative model comparison criterion and to identify the winning model among candidates being tested (i.e. the model with the best trade-off between the quality of the prediction and the complexity, say Model 1 in our example). The preponderant omission of model simulations in the field may betray how common the misconception is that a relative model comparison result is sufficient to conclude that the winning model accounts for a specific “behavior of interest” (i.e. the behavioral difference between A and B in the example; Figure 2). In fact, this is not the case, since the model’s predictive performance and its capacity to reproduce the behavior of interest are formally independent.

To further clarify this point, let us systematically explore and interpret possible relationships between relative model selection criteria and model simulation in our schematic example. We will consider all possible relationships between a given relative model comparison result (i.e. “Model 1 fits better than Model 2”) and the corresponding model simulation results.

First, even the rather extreme situation in which the Model 1 (the winning model) is not capable to reproduce the difference between conditions A and B, whereas Model 2 does, cannot be formally ruled out, uniquely based on a relative criterion (Figure 2A). Although unlikely, this situation can nonetheless occur in cases in which the effect size is particularly small and the model particularly complex, so that the penalization of the extra free parameters cannot be overcome. This rather odd configuration should motivate an amendment in the task design, in order to magnify the effect (more trials or contrast), or the conception of a more parsimonious model, capable of generating the behavior of interest.

Second, it could be possible that neither (Figure 2B) or both (Figure 2C) models surpass the rejection criterion, which therefore do not afford to identify a winning model. This can be easily the case when the behavioral data may be projected into multiple, possibly orthogonal, dimensions (a frequent situation, since behavioral tasks are rarely mono-dimensional). In our example, these dimensions are the average level of responses and the difference between conditions, respectively. As suggested by the relative criterion, in both these intermediate cases Model 1 does account better for the behavioral data, because it generates an overall higher level of responses compared to Model 2. However it happens that these results are completely neutral in respect to our question of interest, which is proposing a theory explaining the difference between A and B. It is nevertheless true that in both cases we could have confidently rejected Model 2, in favor of Model 1, if we were interested in the average level of responses (instead of the difference between A and B). However, in both cases, relative model comparison criteria being blind to the rejection criteria (in our example the difference between A and B), the results do not allow to conclude in favor of any of the two models as the best candidate theory explain the effect of interest.

Finally, it could be that Model 1 proves itself capable of reproducing the A>B contrast, whereas Model 2 does not (Figure 2D). In this case relative model comparison and model simulation analyses concordantly point out the rejection of Model 2 and the acceptance of Model 1 as a possible explanation for such behavioral effect. In this case, if Model 1 and Model 2 have the same number of free parameters, we can ague that the relative model comparison is even useless, since the falsification of Model 2 is sufficient to leave Model 1 as the best candidate theory (among those tested) to explain the effect of interest.

## Relative model comparison has limited explanatory/interpretational power

To summarize, in our simplified example, out of four possible relationships between the same relative model comparison result and different model simulations performance, only one fulfilled the necessary and sufficient conditions to formally conclude in favor of a “winning model”. However, only rarely model-based studies push their analyses that far. This is probably due to the misunderstanding of the complementarity between parsimony and falsification in theory selection. Whereas relative model comparison aims to identify the most probable model among tested candidates, only model simulation can provide *the reasons* of its good (or bad) predictive/fitting performance, thus allowing to conclude on the possible relation about a given model and behavioral phenomenon. In other terms, whereas the relative model comparison criteria inform on “*which mode*l” is the best, the model simulation analyses, by examinating models’ behavior, inform on “*why this model*” is the best. Accordingly, whereas demonstrations uniquely based on relative model comparison criteria are doomed to produce tautological conclusions with no explanatory power in the form of “Model A wins because it has the lower BIC”, only the analysis of model simulation can provide an insight on the behavioral (or neurophysiological) bases of the rejection of a given model.

In summary, given the widespread absence of model simulations to support the results of relative model comparison, it cannot be excluded that some previous computational modeling studies (up to 82%, in our sample) have incorrectly condemned or praised particular computational theories, with no compelling evidence, since only model simulation can provide explanatory power to relative model comparison.

## Model space: does size matters?

When dealing with model comparison a big issue is the selection of the model space. By model space, we mean the set of computational models that are compared in a given study. By definition, there is no theoretical upper limit to the model space – even in simplest paradigms. However, model simulations can be used to define a lower limit of a model space: a model space cannot be considered “sufficient” until there is, at least, one model that is able to reproduce the effect of interest.

Intuitively, it may be argued that the more models are compared, the stronger our model comparison result is. Here, in line with our arguments presented above, we argue that a small, well-justified model space is to be preferred to a larger one. The logic of scientific progress imposes that we replace old theories with new ones, as soon as we are able to show that an old theory cannot account for a new experimental findings^13^. Following from this view, the essential model space should include a “reference” (the model considered as the currently accepted solution for a given computational process) and a “target” model (the new model proposed to replace the “reference” one). Of course, this is quite an idealized case, since, as opposed to physics for example, in cognitive neuroscience there is rarely such a unique “reference” model for a cognitive process. However, on the other side, the invention of new *ad hoc*, often “straw-man”, models, neither “reference” nor “target”, may undermine the clarity and the logic of a computational modeling study.

## Proposed guidelines

In this section, we will propose archetypal analytical guidelines for computational modeling studies of brain function and behavior, which include both relative model comparison and model simulations (see ^17^ for an example of a study including the whole process).

1. Define a task, which implements a cognitive process of interest and include conditions, which are supposed to challenge different ways in which this process is computationally implemented.
2. Simulate *ex ante* the two (or more) competing computational theories across a large range of parameters (sometimes called a ‘parameter recovery’ procedure) in order to ensure that the task allows the discrimination of the two models (i.e. their model predictions diverge in front of a key experimental manipulation). Concomitantly verify that, under the experiment conditions, a given relative model comparison criterion allows to correctly retrieve in both cases the true generative model as the “winning model” (model recovery: see Box 3).
3. Run the experiment and analyze the data to verify that subjects’ behavior was affected by task conditions (model-free analysis).
4. Fit the competing computational models to the data, in order to obtain, for each model an estimation of the best fitting model parameters and an approximation of the model evidence, that trades-off the quality of fit and model complexity.
5. Simulate *ex post* the models, using for each subject the best fitting parameters, in order to verify that only the best fitting model, and ideally not the rival one(s) is capable to reproduce the effect that we are trying to explain.

#### Box 3 model comparison criteria trade sensitivity with conservativeness

The importance of analyzing model simulations is even greater when considering there is no such a thing as a ‘perfect’ model comparison criterion. Most of them are rooted in Bayesian statistics and represent different approximations of the true model evidence. As such, they display different properties in terms of a sensitivity-conservativeness tradeoff. For example, the Bayesian information criterion (BIC)^30^ is typically more conservative, compared to the Akaike Information Criterion (AIC)^31^ or the Laplace approximation of the model evidence^32^ (when the prior over the parameters are relatively flat and non informative as in previous studies^20,33^). Crucially the appropriateness of a given model comparison criterion (given that all are approximations and thus inaccurate) is dependent on the predicted effect size in the experimental data and the models in consideration, with some criteria being more prone to false positive or false negative, depending on the case.

To illustrate this point, we simulated data using an influent reinforcement learning and decision making task^34^. The task is a two-armed bandit game in which options are characterized by reciprocal reward probabilities. The task includes a phase during which the reward contingency is stable and a phase in which it is volatile (i.e. frequently reversed) (Figure Box 3A). It has been robustly showed that humans optimally adjust their learning rate, so that it is lower in the stable compared to the volatile phases. We simulated data in this task with a standard reinforcement-learning model, with no effect of volatility on learning rates (“No modulation” case: α_S_=α_V_) and an adaptive model, with an effect of volatility on learning rates (“Modulation” case: α_S_<α_V_). The data were simulated with as short version of the task, lasting 60 trials, and a long version, lasting 180 trials (as in the original paper) (Figure Box 3B). We then fitted the simulated data by maximum likelihood and LPP maximization (assuming the prior on the learning rate beta(1.1,1.1) and that on the temperatures gamma(1.2,5.) as in previous studies^20,33^) with two computational models (Figure Box 3C). Model 1 has only 2 degrees of freedom and assumes the same learning rate in the stable and the volatile phases. Model 1 is the true generative model for the “No modulation” simulations. Model 2 has 3 degrees of freedom and assumes different learning rates in the stable compared to the volatile phase. Model 2 is the true generative model for the “Modulation” simulations. The two models are nested, so that the likelihood of the data given the model can only increase moving from the simplex to the more complex model. We compared in each simulation type (“No modulation” vs. “Modulation” and “Short task” vs. “Long task”) the two models using the BIC and the LPP: two commonly used quality of fit criteria that trade off likelihood and model complexity. “Correct” inferences are made, when the model comparison criterion “detects” the true generative model, even accounting for its extra complexity and “rejects” the others. “Incorrect” inferences are made when either a more complex model is identified as the best fitting one (a condition known as “overfitting”) or when a less complex model is identified as the best fitting one (a condition that we call “underfitting”) (Figure Box 3C). The results strikingly show that, in our example, the two model comparison criteria may disagree, with one being inevitably in error. More specifically the BIC, more stringent, leads to false incorrect rejection (especially when the task is short, even if the effect of the modulation of the learning rates is quite evident in the simulated behavior) whereas the LPP, more permissive, leads to incorrect detections (Figure Box 3D). This example further highlights that model selection cannot rely only on relative model comparison, and would crucially benefit from looking at model simulations. In our example for instance a quick look to the model simulations would have crucially informed about a model’s capacity to reproduce (or not) quick reversals in the volatile phase. Considering the influence of task’s design (e.g. number of trials) on relative model comparison results, it could be advisable to test *ex ante* the sensitivity of different criteria under different task conditions to optimize the design and identify the most adequate model comparison criterion for a given task and model. Such procedure (that can be defined “model recovery”) would consist in simulating two datasets with two different models and verify (for a given set of models and task specification) which relative model comparison criterion avoid both over- and under-fitting^17^. Alternatively, especially when proposing an original model for a new behavioral task, it would preferable to select the design, such as different relative comparison criteria convergence to the same result^18,20,35^.

**Figure Box 3:**
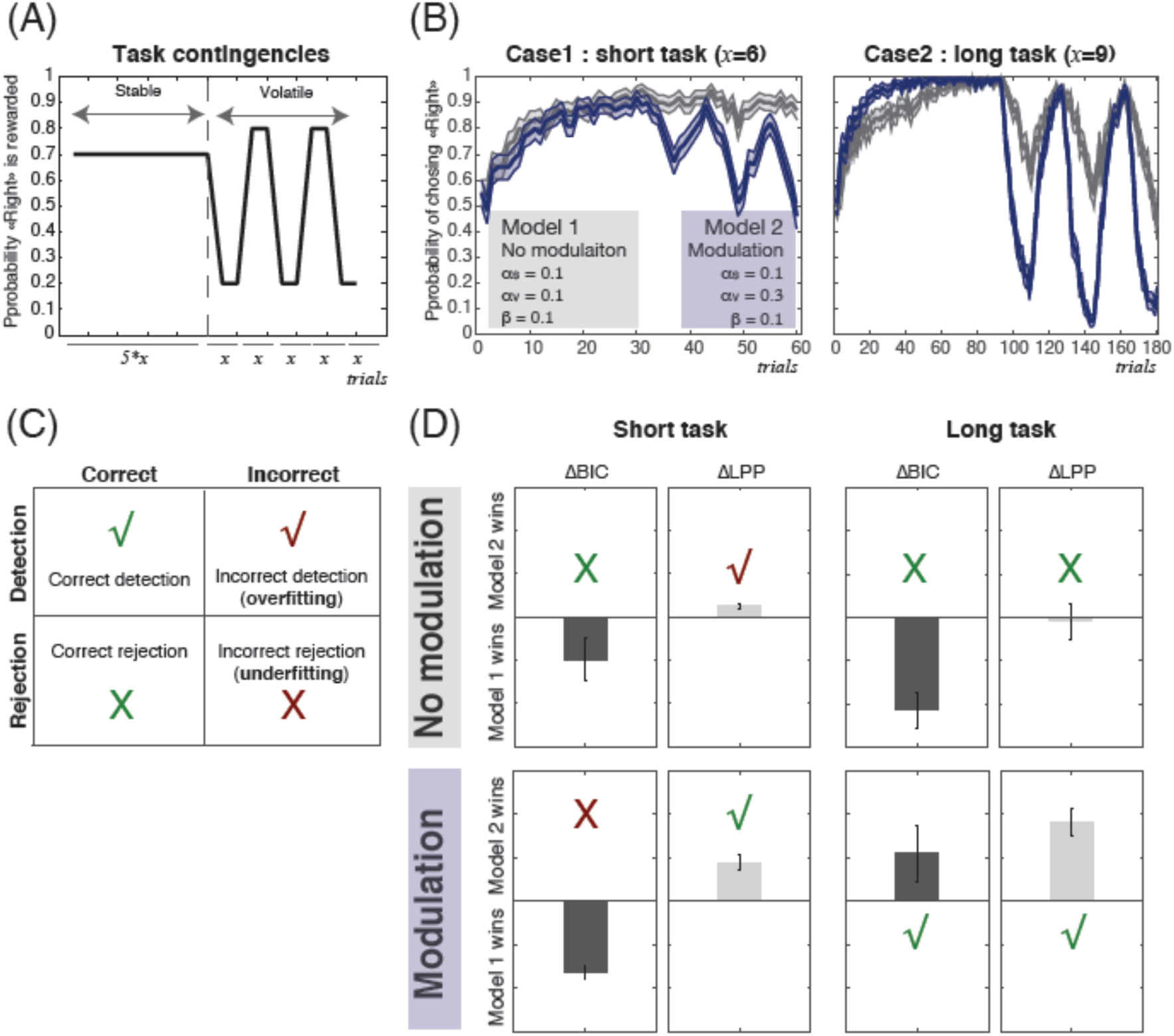
comparison between model comparison criteria. **(A)** Task design. The task includes a stable phase, where the same option is rewarded 70% of the time. The task also includes a volatile phase during which reward contingencies are frequently reversed. The stable and volatile phases last for an equal number of trials, with each volatile episode being five times shorter than the stable phase. (**B**) Simulated data. The learning curves represent data simulated from this task with a standard RL algorithm (in grey; “No modulation” case) and a model that uses a higher learning rate in the volatile compared to the stable phase (in blue; the “Modulation” case). Even in an overall short version of the task (60 trials) choice reversals in the volatile phase are noticeable in the learning curves. **(C)** Expected results and results’ taxonomy. In the “No modulation” case the winning model should be “Model 1” (a simple Q-learning with no modulation of the learning rate) in the “Modulation” case the winning model should be “Model 2” (a model with different learning rates in the stable and volatile phases). Possible results: a correct detection occurs when relative model comparison correctly identifies the more complex model as the “best” model; an “incorrect detection” occurs when relative model comparison wrongly identifies the more complex model as the “best” model (over-fitting); a “correct rejection” occurs when relative model comparison correctly rejects the more complex model as the “best” model; finally an “incorrect rejection” occurs when relative model comparison wrongly rejects the more complex model as the “best” model (under-fitting). **(D)** Model comparison results. The bars represent BIC and LPP differential comparing the reference model (Model 1, two degrees of freedom) with a more complex model (Model 2, three degrees of freedom). The graphs are centered a zero. Positive values indicate that more complex model (Model 2) “wins” compared to the reference model (Model 1).

In practice, computational model-based cognitive neuroscience studies mainly belong to two categories that differ in the place given to the behavioral data and computational models, respectively. The first category of studies, “behavior first”^18,19^, implicates the authors looking for a computational explanation for a previously documented behavioral phenomenon of interest. The second category of studies, “modeling first”^20^, concerns cases in which the aim of the study is to discriminate between competing computational theories. Typical “behavior first” studies may rely only in steps 3-to-5 (since the behavioral task precedes the modeling attempt). On the other side a typical “modeling first” study may not require steps 4-5, if the competing models produce mutually exclusive predictions. Note that the two approaches can be combined. For example the “modeling first” approach may also require *ex post* model simulation, if the two models provide partial overlapping predictions for particular ranges of parameters. Model fitting can be seen as the way to get an insight on the value the parameters could take in the real population (e.g., in the particular case of reinforcement learning, different learning rates such as α=0.2 or α =0.8 do not generate the same learning curves). This is particularly relevant for nested models in which a more complex model can be reduced to a simpler for some specific parameter values.

Note also that these principles and guidelines could be applied to neural data, acknowledging that the simulation of neural data is more complicated, given that the underlying generative process is less understood and may require additional assumptions (but see^21^ for recent evidence challenging the possibility of selecting models based on neural data).

Finally, we would like to specify that we are not arguing that all model-based cognitive neuroscience studies need a model selection procedure (for example when a model has been extensively previously documented or when the model represents an analytical tool and not the object of the study), but that, if a model selection procedure is implemented, it is equally important to investigate both the model’s parsimony, with relative model comparison criteria, and the model’s simulations to understand the reason why the competing model is rejected – and why the winning model has been selected.

## Concluding remarks

In natural sciences, it has been proposed that the epistemological specificity of a computational modeling approach, compared to a model-free one, is that, the latter investigates directly the natural phenomenon of interest, whereas the former builds an artificial representation of the natural system (model) and study its behavior^22^. The ability to reproduce (or not) the natural phenomenon is used to accept (or reject) the model as an accurate representation of the natural system (and its internal laws). In other terms, computational theories have been argued to differ from non-computational, descriptive theories in that the computational theories are capable of generating (simulated) data, which can be then analyzed and compared to empirical data. From this perspective, it clearly appears that relative model comparison results should not be the final outcome of a computational cognitive study, but more likely they should be considered as a quality check, and followed by the comparison of the model simulations of two competing models in respect to a behavioral (or neural) phenomenon of interest. In fact, computational models are not merely supposed to “predict” data, but to mimick the generative processes underlying the behavioral phenomenon of interest. Accordingly, in cognitive neuroscience, computational theories have helped proposing and refining neurocognitive theories via a “reverse engineering” approach, that consists in understanding the brain mechanisms by trying to built algorithms capable to reproduce human behavior^6,7^. After careful consideration of the recent literature, whereas it is uncontestable that computational modeling is taking increasing importance in cognitive sciences (especially in the field of learning and decision making), we also found that the importance of studying the models’ simulations has been overlooked, in favor of relative model comparison techniques. Here we showed, with theoretical argumentation and model simulations (see **Boxes**) that this *lacuna* prevents from realizing the full potential of the computational approach in cognitive neuroscience.

## Acknowledgements

SP was supported by a Marie Sklodowska-Curie Individual European Fellowship (PIEF-GA-2012 Grant 328822). EK is supported by an advanced research grant from the European Research Council (ERC-2009-AdG-250106). VW is supported by a junior researcher grant from the French National Research Agency (ANR-14-CE13-0028). We thank Charles Findling for checking the English and Mehdi Khamassi and Joaquin Navajas for useful comments.

## Footnotes

a “*Pluralitas non est ponenda sine necessitate*.” Plurality is never to be posited without necessity. (*Quaestiones et decisiones in quatuor libros Sententiarum cum centilogio theologico*, Book II) (A.D 1319)

b The full list of the included articles can be obtained by emailing the corresponding author.

